# Plant-parasitic nematode microRNAs hijack plant AGO1 to induce host-cell reprogramming

**DOI:** 10.64898/2026.02.13.705329

**Authors:** Ange Dussutour, Yara Noureddine, Martine da Rocha, Oussama Yahmi, Abile T. Mohammed, Karine Mulet, Pauline Foubert, Ercan Seçkin, An-Po Cheng, Jérôme Zervudacki, Lionel Navarro, Arne Weiberg, Michaël Quentin, Bruno Favery, Stéphanie Jaubert

## Abstract

Cross-kingdom RNA interference (ckRNAi) is emerging as a mode of inter-organismal gene regulation, yet mechanistic examples in plant–metazoan interactions remain limited. Here, we demonstrate miRNA-driven ckRNAi in the nematode–plant pathosystem. Root-knot nematodes are among the most destructive plant pathogens, reprogramming root tissues to develop into galls containing multinucleated, hypermetabolic giant feeding cells essential for parasitism. AGO1-associated small-RNA immunoprecipitation (AGO1-RIP) from tomato galls revealed the selective in planta loading of 10 M. incognita miRNAs into host AGO1. Integrating degradome profiling, target prediction, and dual-luciferase reporter assays, we validated miRNA-directed silencing of nine tomato transcripts by four secreted nematode miRNAs. These targets map to major pathway classes involved in immune signaling, metabolic regulation, and cellular reprogramming linked to feeding-site establishment. Functional analyses further show that the nematode-secreted miR-2b is enhances giant feeding cell development. Comparative AGO1-RIP in Arabidopsis thaliana identified a conserved subset of AGO1-loaded nematode miRNAs, including miR-2b and miR-100, consistent with shared small-RNA effectors across hosts. Finally, the overlap between AGO1-loaded miRNA families and helminth secreted small-RNA repertoires supports evolutionary convergence on RNA-based virulence strategies. Collectively, our findings establish miRNA-mediated ckRNAi as a mechanistic component of plant–root-knot nematode interactions and provide a framework for leveraging RNA-based vulnerabilities for nematode control.

## INTRODUCTION

RNA interference (RNAi) is a highly conserved mechanism of control over gene expression through small non-coding RNAs (sRNAs) (Vaucheret et al., 2004; Bartel, 2009; Bartel, 2018). These sRNAs are typically 18-25 nucleotides (nt) long and are incorporated into Argonaute (AGO)-containing RNA-induced silencing complexes (RISC), which recognize complementary messenger RNAs (mRNAs) by base-pairing and direct posttranscriptional gene silencing (PTGS) by endonucleolytic cleavage and/or translational repression (Ameres and Zamore, 2013). MicroRNAs (miRNAs) constitute a major class of sRNAs in plants. They frequently act through extensive complementarity to guide the silencing of target mRNAs, thereby shaping development, stress adaptation, and immune regulation.

Beyond their canonical intracellular functions, sRNAs have emerged as mobile signals that can act on an organism other than the one in which they were produced, during interactions between these two organisms. Over the last decade, evidence has accumulated that pathogens and symbionts can transfer sRNAs into host cells to reprogram gene expression and promote colonization, persistence, or mutualistic outcomes (Weiberg et al., 2013; Buck et al., 2014). This interorganismal regulation — commonly referred to as cross-kingdom RNA interference (ckRNAi) — has been documented in diverse biological systems (Cai et al., 2019; Qiao et al., 2023). In plants, several fungi, oomycetes, and bacteria have been reported to deploy sRNAs that engage host AGO proteins and suppress host immunity, establishing sRNAs as a distinct class of virulence signals alongside proteins and metabolites (Wong-Bajracharya et al., 2022; Chowdhury et al., 2024; Zhang et al., 2024). Models of the mechanisms underlying sRNA exchange are continually being refined, with the incorporation of roles for extracellular RNA carriers and vesicle-associated pathways, but two prerequisites for functional ckRNAi have consistently been identified: sRNAs must access the appropriate host silencing machinery and must exert sequence-specific regulatory effects on host transcripts. The demonstration of functional engagement is crucial, because non-host sRNAs may be detected in infected tissues due to extracellular contamination or extraction artifacts.

Metazoan–plant ckRNAi has been characterized in much less detail than plant-microbe systems. A few studies have provided evidence that herbivorous insects can deliver regulatory sRNAs that modulate plant gene expression, but the extent to which this phenomenon is generalized among metazoan plant pathogens remains unclear (Zhang et al., 2024; Han et al., 2025). This constitutes a major gap in our knowledge, given the considerable reliance of several metazoan parasites on intimate, long-term interactions with plant tissues, potentially involving RNA-based effector strategies. Determining whether metazoan plant pathogens deploy sRNAs as *bona fide* effectors — *i.e.*, sRNAs that enter host cells, load host AGO complexes and direct the silencing of host targets — would therefore advance our fundamental understanding of host manipulation, potentially expanding the conceptual framework for RNA-based disease control.

Root-knot nematodes (RKNs) of the genus *Meloidogyne* are highly adapted, economically important plant parasites that rank among the most damaging plant pathogens worldwide. *Meloidogyne incognita* is particularly notable for its exceptionally broad host range, infecting more than 5,000 plant species, including most cultivated vegetable crops (Castagnone-Sereno, 2006; Blok et al., 2008; Abad and Williamson, 2010). Despite this diversity of hosts, RKNs reproducibly induce morphologically and functionally similar feeding sites, highlighting their capacity to exploit cellular programs that are conserved in plants. After invading the roots, juveniles establish a permanent feeding site by reprogramming a small number of vascular parenchyma cells into multinucleated, highly metabolically active giant cells that function as strong nutrient sinks required for nematode growth and reproduction. The formation and maintenance of these feeding structures requires extensive remodeling of host transcription and physiology in response to nematode-derived signals (Favery et al., 2020). The esophageal glands of RKNs produce a large repertoire of secreted effectors, which are delivered via the stylet, making it possible to manipulate plant immunity, cell cycle control, development, and RNA metabolism directly (Mejias et al., 2019; Mejias et al., 2022; Rutter et al., 2022). Protein effectors play a well-established role as central drivers of parasitism, but the extent to which sRNAs contribute to nematode-induced host reprogramming remains unclear. Plant-nematode interfaces are already known to allow biologically meaningful RNA transfer in the opposite direction: host-induced gene silencing (HIGS), in which plants express dsRNA/siRNAs targeting nematode genes, can reduce nematode parasitism (Gheysen and Vanholme, 2007).

In this context, miRNA-based ckRNAi is a plausible and potentially powerful strategy. Many core regulatory and signaling nodes are conserved across plant species and miRNA effectors targeting these nodes may explain how a single nematode species can manipulate host pathways common to diverse plants. Moreover, the functional engagement of nematode miRNAs by the plant silencing machinery would provide a direct route for the suppression of host defenses or fine-tuning of developmental and metabolic programs associated with feeding-site formation. However, stringent evidence is required to support claims of ckRNAi in complex infected tissues, as galls contain a mixture of plant and nematode material and RNA profiles can be confounded by tissue admixture or non-specific associations during extraction. Approaches in which functional loading into host AGO proteins is used as the readout, rather than the bulk detection of parasite RNAs, therefore provide a stringent framework for distinguishing between ckRNAi candidates and background.

We investigated the engagement of *M. incognita*-derived miRNAs by the plant AGO1 pathway during gall development. We used AGO1-associated sRNA immunoprecipitation (AGO1-RIP) from infected tomato roots as a functional readout for the sRNAs present in the host RISC, coupled with stringent controls to identify nematode sRNAs potentially displaying non-specific association with AGO1 during sample processing. We then used degradome evidence *and in planta* reporter-based assays to investigate whether candidate nematode miRNAs directed the sequence-specific cleavage of host transcripts. By performing AGO1-associated profiling on phylogenetically distant host plants, we investigated whether a subset of nematode miRNAs was widely deployed across hosts and whether their predicted/validated target engagement was consistent with the modulation of conserved pathways linked to feeding-site formation. Finaly, functional analyses further establish nematode-secreted miR-2b as a determinant of giant feeding cell formation. Together, these findings establish a functional framework for evaluating miRNA-based ckRNAi in plant–nematode interactions and clarifying the role of nematode miRNAs as effectors in the molecular dialog underlying RKN parasitism.

## RESULTS

### Genome-wide identification of *Meloidogyne incognita* miRNAs

We characterized the *M. incognita* sRNA repertoire during parasitism by generating sRNA libraries from freshly hatched, preparasitic second-stage juveniles (J2) and from parasitic stages embedded in tomato galls at 7 and 14 days post infection (dpi). High-quality reads were aligned with *M. incognita* genome assembly V3 (Blanc-Mathieu et al., 2017) and used for *de novo* miRNA annotation.

By integrating miRDeep2 and ShortStack predictions and curating conserved loci, we identified 288 *MIR* genes in the *M. incognita* V3 genome, including 119 miRNAs with orthologs in miRbase. These 288 *MIR* genes were grouped into 158 miRNA families (Supplemental Figure 1 and Supplemental Table 1) and organized into 17 genomic clusters (Supplemental Table 2). A substantial proportion of these *MIR* loci corresponded to putative lineage-specific miRNAs (i.e., with no detectable homologs in available nematode or plant genome sequences; number = 167). A core set of 15 conserved miRNAs (miR-100, miR-1, miR-71, lin-4, miR-5359, miR-92, miR-124, miR-50, miR-86, miR-87, miR-7904, miR-2a, miR-2b, miR-279, miR-252; ranked by abundance), and one *Meloidogyne-*specific miRNA (miR-minc70) dominated the sRNA pool (Figure 1; Supplemental Table 3). A small number of conserved miRNA families therefore account for a large proportion of total miRNA reads despite extensive lineage-specific diversification.

**Figure 1.**
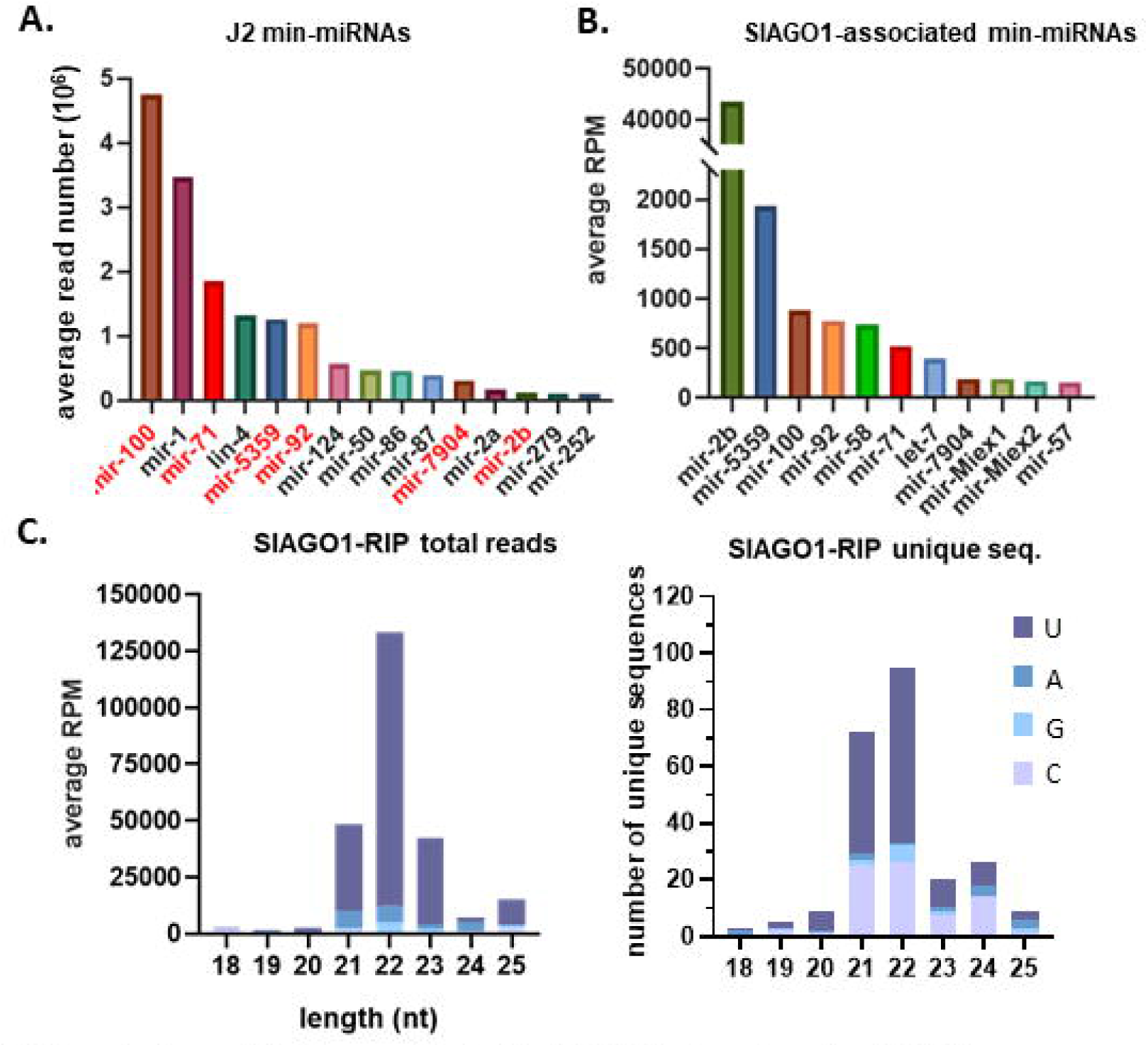
Nematode small RNAs associated with SlAGO1 in tomato galls at 14 dpi. (A) Top 15 most abundant *M. incognita* miRNAs in sRNA-seq libraries from pre-parasitic J2s, shown as mean abundance across three biological replicates in read per million (RPM). (B) *M. incognita* miRNAs detected in association with tomato AGO1 (SlAGO1) in galls at 14 dpi. miRNAs highlighted in red correspond to min-miRNAs detected in association with *S. lycopersicum* AGO1 in galls. (C) SlAGO1-RIP size and 5′-nucleotide profiles from two biological replicates of tomato galls, shown as average (RPM) and number of unique sequences for each sRNA size class.

### *M. incognita* miRNAs hijack host AGO1 during infection

RKN are obligatory sedentary endoparasites, which makes it difficult to isolate the secretion they produce *in planta*. We investigated whether *M. incognita* sRNAs were delivered to plant cells and engaged the host RISC, by performing AGO1 RNA immunoprecipitation (AGO1-RIP) on tomato (*Solanum lycopersicum*) root galls at 14 dpi with anti-AGO1 antibodies previously validated for plant AGO1-RIP assays (Dunker et al., 2021). Galls are a mix of plant and nematode tissues. We therefore used a stringent negative control in which uninfected tomato roots were mixed with preparasitic J2s immediately before tissue disruption, thereby enabling to identify nematode sRNAs that might bind AGO1 non-specifically during extraction.

We removed ribosomal RNAs (rRNAs), transfer RNAs (tRNAs), small nuclear RNAs (snRNAs), and small nucleolar RNAs (snoRNAs) reads, retaining only unique sRNA sequences present at a frequency of >50 reads per million (rpm) in at least two biological replicates. We identified 236 unique sRNA sequences associated with SlAGO1 displaying the expected enrichment in 22-nt species with a 5′ uracil residue (Figure 1, Supplemental Table 4). The mapping of AGO1-associated sRNAs onto the tomato (ITAG 3.2) and *M. incognita* (V3) genome sequences resulted in 218 sequences being assigned to tomato and 18 to *M. incognita*.

Eleven of the 18 nematode-derived AGO1-associated sRNAs were annotated *M. incognita* miRNAs (Table 1). These 11 nematode miRNAs accounted for 19.6% of all AGO1-RIP reads in infected galls, corresponding to an enrichment of about 50-fold relative to the negative control, in which nematode sRNAs accounted for 0.4% of AGO1-associated reads. The set of AGO1-associated nematode miRNAs consisted of nine conserved miRNAs (miR-2b, miR-5359, miR-100, miR-92, miR-58, miR-71, let-7, miR-7904, miR-57; ordered by abundance) and two *Meloidogyne*-specific miRNAs (miR-Miex1 and miR-Miex2).

**Table 1:**
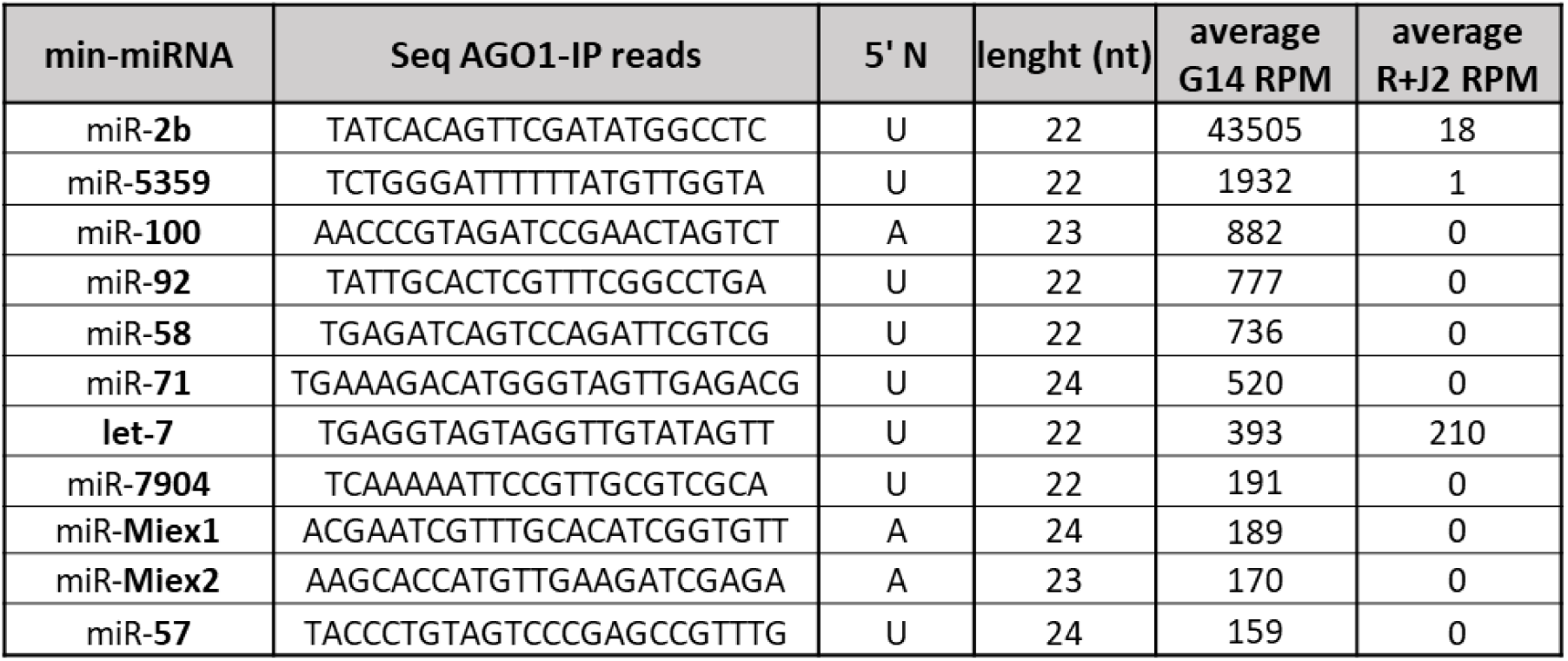
List of SlAGO1-associated *M. incognita* miRNAs in tomato galls at 14dpi.

Three nematode microRNAs — miR-2b, miR-5359 and let-7 — were also found in the control. An analysis of differential expression between galls and the negative control revealed a massive enrichment in miR-2b and miR-5359 in galls, whereas the abundance of let-7 was similar in galls and the control (Supplemental Table 5). This result suggests that the association of let-7 with SlAGO1 may be due to non-specific association during the grinding together of plant and nematode tissues. We therefore excluded let-7 from further functional analysis. Overall, these results demonstrate a strong and specific enrichment of a defined set of *M. incognita* miRNAs in the SlAGO1 complex during infection, supporting their active delivery in host cells.

### The selective enrichment of AGO1-associated nematode miRNAs does not correlate to nematode miRNA abundance

We then investigated whether the abundances of nematode miRNAs in SlAGO1-RIP reflected their overall abundance in the nematode. In tomato gall AGO1-RIP datasets, miR-2b was the most abundant nematode miRNA species — at a median of 622 rpm among nematode-associated reads and with a mean accumulation of ∼43,000 rpm overall — accounting for 14.7% of all AGO1-associated sRNAs and constituting the dominant component of the nematode-derived fraction. At whole-library level, miR-2b was the second most abundant AGO1-associated sRNA species, surprisingly exceeded only by the sly-miR-482b miRNA of the host plant, tomato (Supplemental Table 4).

We then assessed the abundance of *M. incognita* miRNA in sRNA-seq datasets derived from preparasitic J2s (Figure 1 and Supplemental table 3). Several nematode miRNAs detected in SlAGO1-RIP (miR-100, miR-71, miR-5359, miR-92, miR-7904, and miR-2b) were among the top 15 most abundant miRNAs in preparasitic stages. By contrast, other SlAGO1-associated miRNAs (miR-Miex1, miR-Miex2, miR-58, miR-57) were not among the most abundant miRNAs in nematode bulk datasets. Conversely, several nematode miRNAs found to be highly expressed in J2/G14 datasets (miR-1, miR-124, lin-4, miR-50, miR-87, miR-279) were absent from SlAGO1-RIP libraries. The ranking of AGO1-associated nematode miRNAs by abundance therefore displayed limited concordance with the overall abundance of these molecules in nematode sRNA-seq datasets. An extreme example was provided by miR-2b, which had a low abundance in nematode bulk datasets but was very frequent among AGO1-associated reads *in planta*. Together, these results indicate that the nematode sRNAs delivered to plant cells and loaded on plant AGO1 constitute a selective subset of nematode miRNAs rather than a passive reflection of the overall nematode miRNA pool.

### Identification of the targets in tomato of the microRNAs secreted by *M. incognita*

For the identification of putative tomato transcripts targeted by AGO1-associated nematode miRNAs, we combined degradome analyses with computational target prediction. We first reanalyzed a tomato 14 dpi gall degradome library (Noureddine et al., 2023) with Cleaveland4 and the 10 nematode miRNAs retained after the exclusion of let-7. Using ≤2 as the degradome category cutoff, we identified 63 candidate tomato targets of the 10 nematode miRNAs (Supplemental Table 6). Target numbers were heterogeneously distributed between miRNAs: miR-2b, miR-5359, miR-92, and miR-Miex1 were associated with 43 targets, with miR-5359 associated with the largest number of targets for any single miRNA (22 targets). By contrast, miR-58, miR-Miex2, miR-7904, miR-57, and miR-100 together accounted for only 13 targets, and no degradome-supported target was recovered for miR-71 with the application of these criteria.

Degradome sensitivity can be reduced by tissue complexity, nematode sRNA dilution and the rapid degradation of cleavage products. We therefore performed complementary *in silico* prediction with psRNATarget. This analysis identified 201 putative targets (mean of ∼20 targets per miRNA; Supplemental Table 7). Consistent with the degradome analysis, the number of predicted targets was highest for miR-5359 (57) and lowest for miR-57 (2). Relative to degradome-based results, psRNATarget predicted larger target sets for miR-7904, miR-58, miR-Miex2, and miR-71 (25-31 predictions per miRNA). Nineteen transcripts were predicted as target by both analyses for eight of the secreted nematode miRNAs. 29 candidate targets were then selected for further experimental testing of silencing *in planta* (Supplemental Table 8).

### *In planta* validation of nematode miRNA silencing activity with dual-luciferase reporters

We investigated the ability of nematode miRNAs to silence predicted tomato targets in plant cells, in an *in planta* dual-luciferase reporter (DLR) assay (Liu et al., 2014) and by the expression of artificial miRNAs (Schwab et al., 2010). For each nematode miRNA, we included a positive silencing control reporter bearing a perfectly complementary target site to ensure expression of the nematode miRNA *in planta* and its efficient silencing. Empty vector was used as a negative cleavage control. The R-luc/F-luc ratios of targets fused to the F-luc 3′UTR and co-expressed with the miRNAs were normalized against the basal R-luc/F-luc ratio of the negative control.

For the nine secreted nematode miRNAs tested, silencing activity was confirmed for miR-2b, miR-5359, miR-92, miR-57, miR-58, miR-Miex1, and miR-7904, which yielded significant decreases in reporter activity for the perfect-complementarity control (Supplemental Figure 2). Silencing strength varied between the miRNAs, with miR-2b and miR-7904 decreasing the reporter activity of the positive control relative to the negative control by ∼65%, whereas miR-92, miR-Miex1, miR-57, and miR-58 yielded more moderate decreases in reporter activity (∼20–45%). The strongest activity was observed for miR-5359, which decreased the F-luc signal of the positive control by >97%. By contrast, no decrease in reporter activity was observed for miR-100 or miR-Miex2 with either of the positive control constructs. These miRNAs were therefore either not expressed or not active in this assay configuration under our experimental conditions.

Nine of the 29 candidate targets in tomato tested displayed significant decreases in F-luc activity following co-expression with the corresponding nematode miRNA. We identified three targets for miR-2b (*Solyc06g067890, Solyc09g091540, Solyc12g049320*), two for miR-5359 (*Solyc11g006760, Solyc11g045110*), two for miR-Miex1 (*Solyc06g072690, Solyc07g065890*), and two for miR-7904 (*Solyc03g118010, Solyc11g005450*) (Figure 2). Thus, four of the nine miRNAs tested silenced nine of the 29 targets *in vivo* (∼31%), providing evidence in support of the functional targeting of specific tomato transcripts by plant AGO1-associated nematode miRNAs (Table 2).

**Figure 2.**
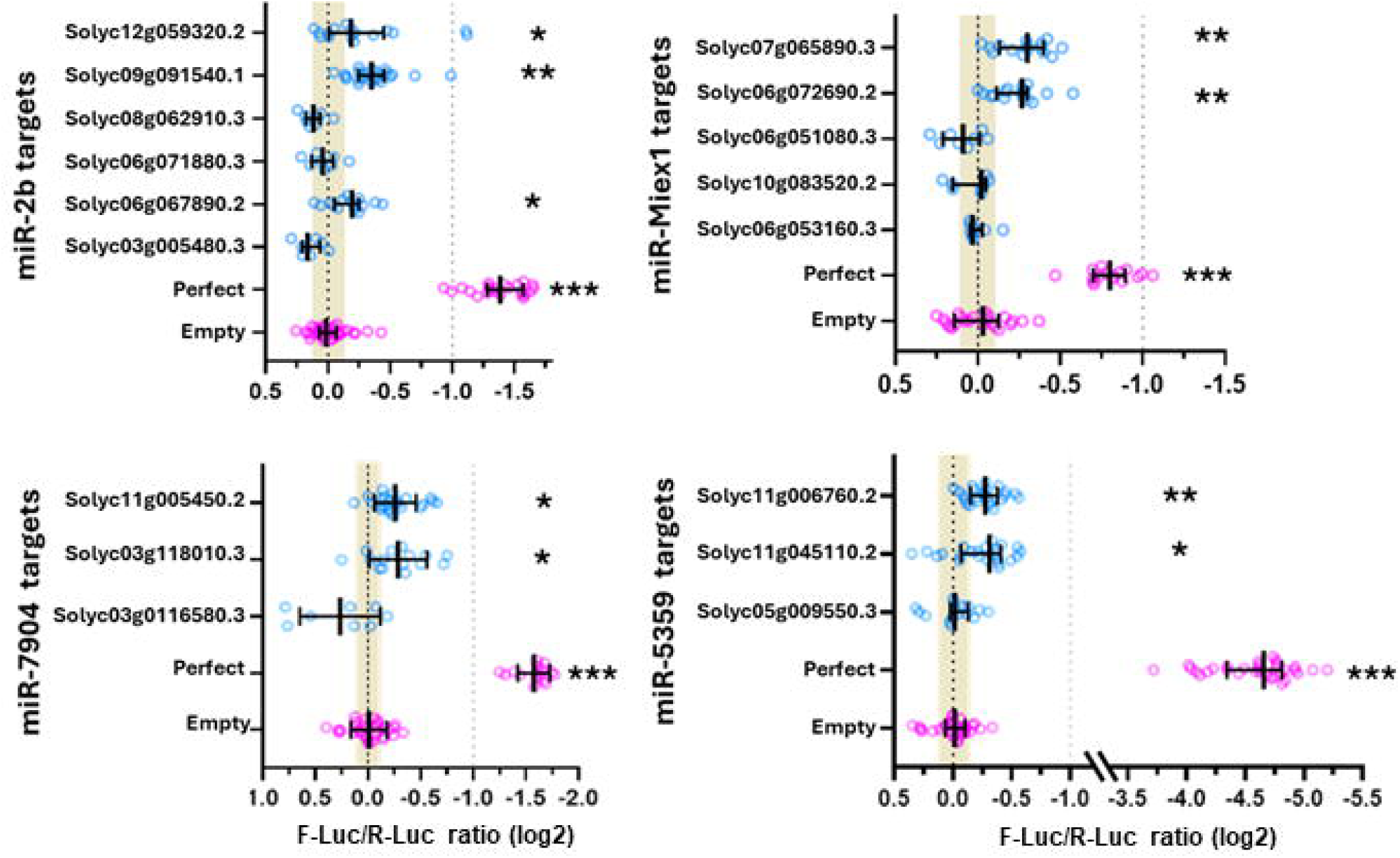
Experimental validation of nematode miRNA–directed silencing of predicted tomato targets. Nematode miRNAs and their predicted tomato target sites were co-expressed in *Nicotiana benthamiana* leaves to assess silencing activity. Target sites were fused to the firefly luciferase coding sequence in a dual-luciferase reporter (DLR) system, with constitutively expressed Renilla luciferase as an internal control. The log2-transformed Firefly/Renilla (F-luc/R-luc) ratio measured in co-infiltrated leaves was normalized to the negative control. A perfectly complementary target sequence (“Perfect”) served as a positive silencing control. For negative-control assays, a “Spacer” corresponding to the native 5′ UTR sequence upstream of firefly luciferase (with little or no complementarity to the tested miRNA) was used. Asterisks indicate statistical significance (unpaired Student’s *t*-test): *P* < 0.05, \**P* < 0.005, \**P < 0.0001*.

**Table 2:**
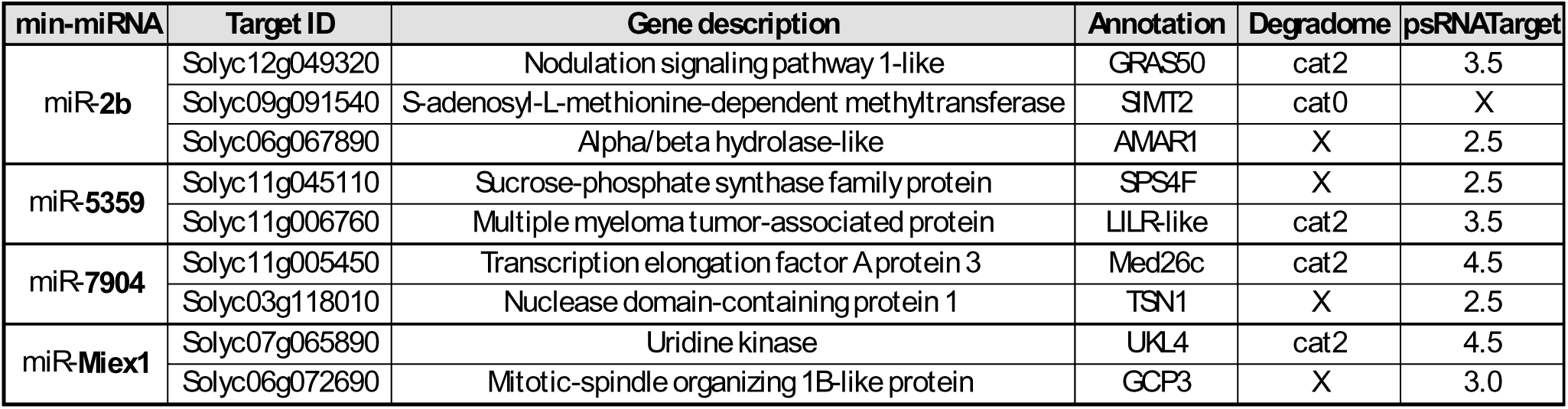
List of targets validated in DLR with gene ID, annotation and prediction method.

### Comparative AGO1-RIP reveals a conserved subset of nematode miRNAs in host plants

Given the broad host range of *M. incognita*, we investigated the possible conservation of the AGO1-associated nematode miRNA repertoire between evolutionarily distant hosts. We performed AGO1-RIP on *Arabidopsis thaliana* galls at 14 dpi and profiled AGO1-associated sRNAs. This analysis detected miR-2b, miR-100, let-7 and miR-57 as nematode-derived miRNAs present in AGO1-associated sRNA pools in *Arabidopsis,* as they are in tomato (Supplemental Table 9), indicating that at least a subset of AGO1-associated *M. incognita* miRNAs are common to divergent host species.

By contrast to the results obtained for miR-2b, we were unable to validate the silencing of the tested putative targets of miR-57 *in planta* and no silencing activity could be assessed for miR-100 in our experimental conditions. We then further assessed conservation of validated target sites of miR-2b for which orthologs were available. The *Arabidopsis* ortholog of the tomato miR-2b target SlMT2 *SALICYLIC ACID METHYLTRANSFERASE-2* (*Solyc09g091540*), *JASMONIC ACID CARBOXYL METHYLTRANSFERASE (AtJMT)* (*AT1g19640*), retained a conserved target site (Supplemental Figure 3). By contrast, orthologs corresponding to other tomato miR-2b targets displayed sequence divergences that decreased the near-perfect complementarity at the predicted target sites, consistent with potential host-dependent differences in cleavage compatibility. Conservation of the miR-2b target site across orthologs points to a conserved miRNA-controlled pathway that likely underpins a core M. incognita infection strategy across its broad host range.

### Conservation and genomic organization of secreted miRNAs in different *Meloidogyne* species

In the *M. incognita* genome, the miRNAs released into the host belonged to 10 genomic clusters (Supplemental Table 2). Clustered miRNAs are transcribed as a single primary transcript (pri-miRNA), which is then processed into multiple precursor and mature miRNAs. Several of the loci encoding miRNA released into the host (including copies of miR-2b, miR-71, and miR-5359) clustered with *MIR* genes encoding miRNAs not detected in AGO1-RIP datasets.

We evaluated the conservation of secreted miRNAs across RKNs, by performing microsynteny analyses to identify precursor (pre-miRNA) orthologs in three additional *Meloidogyne* genomes (*M. enterolobii, M. arenaria,* and *M. javanica*) (Supplemental Table 10). We were able to identify orthologous precursor sequences for most of the secreted miRNAs, generally with similar copy numbers across species consistent with reported ploidy differences. For many MIR *loci*, the surrounding genomic context was also conserved, supporting orthology at locus level.

### The miR-2 family in *Meloidogyne*: copy number, variants, and secretion-associated patterns

The most abundant nematode-derived miRNA associated with host AGO1 was miR-2b, accounting for 86% of all reads assigned to secreted nematode miRNAs (Table 1). We investigated the genomic basis of this enrichment in *Meloidogyne*. In *M. incognita*, we identified eight *MIR-2* loci encoding three mature variants: the AGO1-associated miR-2b and two variants not detectably associated with host AGO1 under our conditions (miR-2a and miR-2c). These variants had identical seed sequences but differed outside the seed region (Supplementary Figure 4).

Small-RNA profiling across developmental stages revealed broadly similar expression patterns for miR-2 variants in preparasitic J2s (Supplemental Table 3). However, despite the detectable expression of multiple miR-2 variants, only miR-2b displayed strong and reproducible enrichment in plant AGO1-RIP libraries, consistent with selective association with the host silencing machinery.

Consistent with our observations in *M. incognita*, miR-2 family members have been detected in the secreted/extracellular fractions of several animal-parasitic nematodes (Sotillo et al., 2020; Chowdhury et al., 2024), indicating that host exposure to miR-2 is observed across phylogenetically distant parasitic systems. We then assessed the genomic organization of miR-2 loci. In many metazoans, *MIR2* genes occur in family-specific clusters, consistent with tandem duplication. By contrast, in nematodes, miR-2 loci are frequently found as singletons or within heterogeneous miRNA clusters. Consistent with this pattern, *M. incognita MIR2* genes were distributed heterogeneously between singletons and heterogeneous clusters, rather than forming a single family-specific cluster (Supplemental Figure 4). The extension of this analysis to additional nematode genomes revealed substantial variation in miR-2 copy number (range: 3 to 11 loci; Supplemental Figure 4). With eight loci, *M. incognita* was one of the species with the highest miR-2 copy number in our dataset, together with *A. suum* (11 loci) and *B. malayi* (10 loci). In species encoding multiple mature miR-2 variants, only a subset of these variants were detected in host-exposed/AGO1-associated datasets, consistent with the variant-specific enrichment observed for miR-2 family members during parasitism.

### miR-2b Promotes Feeding Cell Expansion

We investigated whether loading on the host AGO1 reflected a function in parasitism by generating tomato roots overexpressing either miR-2b or miR-57 using *Agrobacterium rhizogenes* mediated transformation system. We investigated the effect of overexpressing secreted nematode miRNAs within these transformed roots on the formation of feeding cells by measuring the size of the nematode-induced feeding cells within these roots. Tomato roots transformed with an empty vector were used as a negative control. The overexpression of miR-2b in tomato roots induced a clear increase in feeding site size, whereas no effect was observed for miR-57 (Figure 3). Altogether, these findings support a role for miR-2b in promoting feeding cell formation and/or sustaining their maintenance during infection.

**Figure 3.**
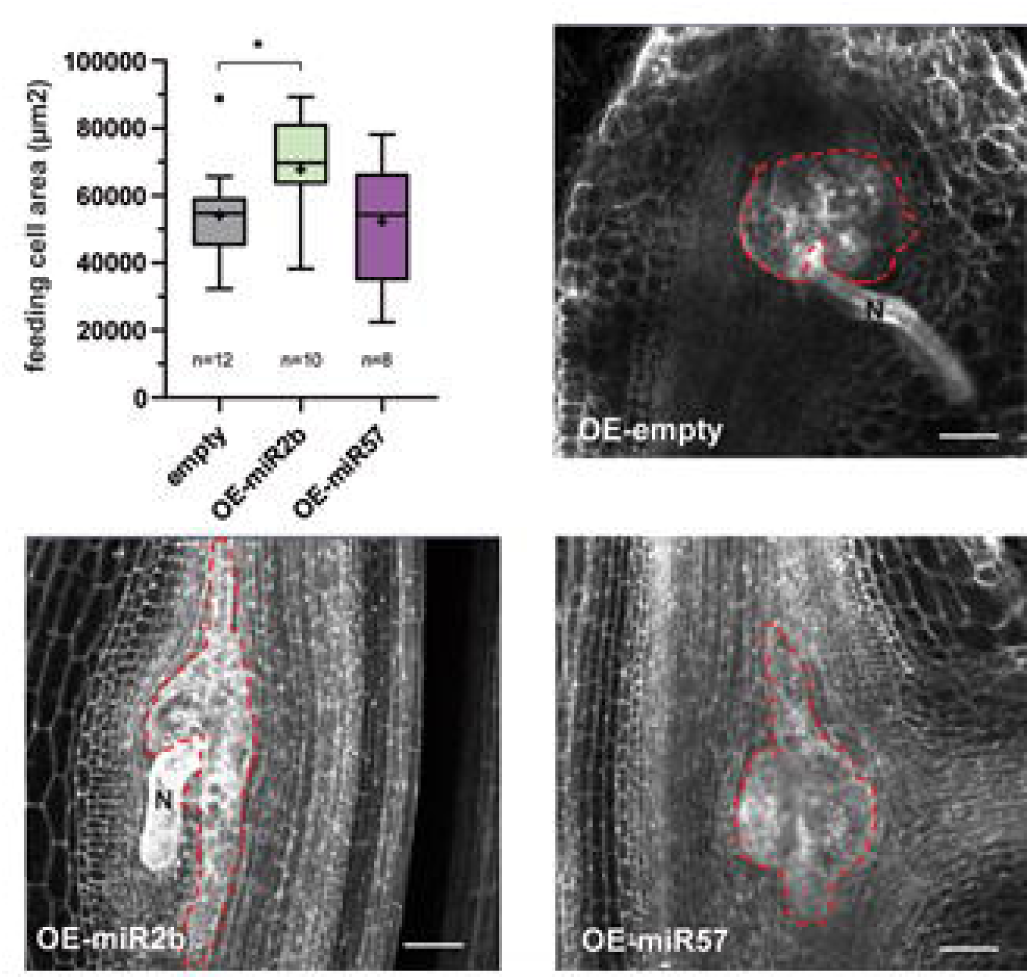
*In planta* overexpression of min-miR-2b increases feeding-site size in tomato galls. The effects of min-miR-2b and min-miR-57 on giant feeding cell development were assessed by quantifying the feeding-site area (giant-cell region; outlined in red) in tomato roots transiently expressing miR-2b or miR-57, compared with empty-vector controls. Galls were collected 2 weeks after *in vitro* infection and cleared using the BABB method (Cabrera et al., 2018) to measure the area (µm²) corresponding to giant cells. Differences were evaluated using Mann–Whitney tests (*P* < 0.025). Box plots show the interquartile range (25th–75th percentiles) with the median indicated by the center line; whiskers represent the minimum and maximum non-outlier values, and points denote outliers. *n* indicates the number of galls analyzed per line and condition. Scale bar, 100 µm. N, nematode.

## DISCUSSION

Our data support a model in which M. incognita deploys a highly selective, miRNA-based ckRNAi strategy: a restricted set of nematode miRNAs enters host root cells, loads into plant AGO1, and hack plant silencing pathway. Functional validation indicates that these miRNAs converge on key host hubs controlling immunity, metabolism, and cell fate, consistent with a coordinated reprogramming of gall development and feeding-site function during infection.

### Selective sorting and AGO1 loading of nematode miRNAs in host cells

CkRNAi is based on the selective transfer of a defined subset of sRNAs into interacting organisms, in which these molecules engage the RNAi machinery and mediate the silencing of target genes (Weiberg et al., 2013; Dunker et al., 2020). Our results are consistent with the effective selection of certain miRNAs for secretion into host cells. The nematode miRNAs associated with the plant AGO1 are not simply the miRNAs most abundant in the nematode. Several highly expressed nematode miRNAs are absent from the host AGO1, whereas some of the miRNAs associated with AGO1 are not among the predominant species in the nematode. This lack of coupling strongly argues that there is selective sorting and/or selective stabilization of a restricted subset of miRNAs for delivery in the host. Moreover, the pronounced enrichment (30-fold) in *M. incognita* miRNAs relative to the negative control displayed by the host AGO1 demonstrates not only the secretion and delivery of these miRNAs into host cells but also their functional integration into the central effector of the plant RNA silencing pathway. Finally, some miRNAs destined for secretion that are encoded by genes clustered with non-secreted miRNA genes within the nematode genome were found to be secreted and loaded into the host AGO1. This organization may suggest that the clustered miRNAs are transcribed in the same cells, implying that delivery to the host is unlikely to be solely due to expression in a specific cell type. However, differential maturation, the stability of mature miRNAs, and posttranscriptional regulation may result in imbalances between the expression profiles of members of the same cluster (Martinez et al., 2008; Jovelin and Cutter, 2014).

### Nematode-secreted miRNAs actively cleave specific tomato transcripts

By combining degradome profiling, *in silico* target prediction, and in planta DLR assays, we identified secreted nematode miRNAs capable of cleaving specific tomato transcripts. Because galls comprise only a few giant feeding cells embedded within proliferating surrounding tissues, cleavage events are expected to occur in a very limited cell population, thereby diluting the signal in whole-gall degradome datasets and likely accounting for the low degradome categories observed for candidate targets. DLR assays revealed modest but reproducible repression (typically ∼10–20%), lower than in some reported plant–pathogen systems (Meng et al., 2021; Mei et al., 2025), and variable across miRNA–target pairs. The absence of activity for miR-100 and miR-Miex2 in our positive control context may reflect technical constraints—such as inefficient amiRNA processing or expression, or suboptimal AGO1 loading—rather than a lack of intrinsic targeting potential.

### Conservation across hosts and implications for host range

*M. incognita* miR-2b, miR-57 and miR-100 were found to be loaded into AGO1 in galls from *S. lycopersicum* and *A. thaliana*, demonstrating the secretion of a small set of conserved *M. incognita* miRNAs in phylogenetically distant host species and suggesting that these miRNAs may be factors involved in the manipulation of conserved plant mechanisms during RKN infection. As in tomato, miR-2b is one of the main miRNAs secreted in *Arabidopsis* galls, supporting the hypothesis that miR-2b plays an important role in *Meloidogyne* infection processes. Nevertheless, the ubiquitous secretion of large amounts of miR-2b and the low level of cleavage site variation between plant jasmonate methyltransferase homologs support the hypothesis that the JA hormonal defense pathway is a major and conserved target in diverse plants within the broad host range of *M. incognita*. Target sequence divergence probably also restricts some silencing activities to specific hosts, potentially contributing to quantitative differences in susceptibility.

### Host pathways targeted by secreted miRNAs

Nine secreted nematode miRNA–tomato gene pairs were validated by DLR assays. The targets of the four active miRNAs (miR-2b, miR-7904, miR-Miex1, and miR-5359) belong to a restricted set of pathways playing key roles in plant-RKN interactions, including host defense processes commonly targeted by ckRNAi in other plant-pathogen systems (Weiberg et al., 2015; Cai et al., 2019) and metabolic and developmental programs particularly relevant to RKN parasitism (Favery et al., 2020). Hormone-mediated immunity (SA/JA/ET) emerged as a prominent node, as silencing was confirmed for SlMT2 (*Solyc09g091540*; targeted by miR-2b), SlMED26c encoding a Mediator complex subunit (*Solyc11g005450*; targeted by miR-7904), the uridine kinase UKL3 (*Solyc07g065890*; targeted by miR-Miex1), and possibly a GRAS transcription factor (*Solyc12g049320*); together, these findings are consistent with an attenuation of defense signaling during infection (Wang et al., 2019; Kong et al., 2020; Cheng et al., 2025). Other secreted miRNA target pathways linked to feeding-site establishment and maintenance, notably energy metabolism and cellular reorganization, were also identified: TSN (*TUDOR-SN nuclease*, *Solyc03g118010*) and SPS4F (a sucrose-phosphate synthase, *Solyc11g045110*), targeted by miR-7904 and miR-5359 respectively, are linked to SNF1-related protein kinase (SnRK1)-dependent metabolic control, a central regulatory hub and master metabolic regulator (Nukarinen et al., 2016; Van Leene et al., 2022). Furthermore, miR-2b and miR-5359 also silence transcripts encoding an alpha/beta hydrolase (*Solyc06g067890*) and a multiple myeloma tumor-associated protein 2 (*Solyc11g006760*), the homologs of which in *Arabidopsis AT1g80280* (*XOPAM-ACTIVATED RESISTANCE 1, AMAR1*) and *AT3G52220* (a LEUKOCYTE IMMUNOGLOBULIN-LIKE RECEPTOR (LILR)-like domain protein) are linked to immune-related functions (Xie *et al*. 2023). Beyond direct defense and metabolic outputs, silencing of the GRAS transcription factor, MED26c, and TSN would be expected to have broader effects on transcriptional and posttranscriptional regulation, given the reported roles of these molecules in symbiotic programs, RNA polymerase II-mediated transcriptional reprogramming, and the RISC-associated control of endogenous miRNA abundance (Jaiswal et al., 2022; Freytes et al., 2024; Jiang et al., 2025). Finally, the targeting of the tomato *GAMMA-TUBULIN COMPLEX COMPONENT* 3 (*GCP3*) homolog (*Solyc06g072690*) by miR-ex1 suggests a role in regulation of cytoskeletal organization and cell division required for formation of in giant feeding cells (Caillaud et al., 2008; Banora et al., 2011). Together, these results support a model in which nematode-secreted miRNAs orchestrate a coordinated suite of regulatory effects that damp down host immunity while directing changes in metabolism and cellular architecture to promote giant-cell development and sustain feeding-site function throughout infection.

### miR-2 family expansion and subfunctionalization

The exceptional accumulation of miR-2b within AGO1 complexes, accounting for a substantial fraction of total AGO1-associated sRNAs, identifies this miRNA as a major cross-kingdom silencing effector operating during infection. The miR-2 family is an ideal framework in which to investigate how miRNA gene expansion can be harnessed for parasitism. *Meloidogyne* species have multiple miR-2 loci encoding different mature variants (miR-2a/2b/2c) with identical seeds, but with only one variant (miR-2b) displaying consistent enrichment among host AGO1-bound sRNAs and ranking among the most abundant nematode-derived sRNAs in infected tissues. This marked asymmetry fits a subfunctionalization model, with miR-2 paralogs partitioning roles between endogenous regulation and cross-kingdom delivery, mirroring patterns described for duplicated protein-coding genes (Blanc Matthieu et al. 2017). the context of the plant RNAi machinery, seed identity alone is insufficient to predict functional equivalence, as base-pairing outside the seed contributes to target recognition in plants, in which near-perfect complementarity is a major determinant of silencing activity (Liu et al., 2014). Accordingly, modest sequence differences between miR-2 variants might dramatically alter the availability of quasi-complementary target sites in host transcripts, while simultaneously affecting secretion-associated sorting. Notably, the poor correlation between abundance and association with the host AGO1 observed for nematode miR-2b indicates that secretion is not driven solely by expression level, providing additional support for the existence of selective trafficking mechanisms.

### Evolutionary convergence of secreted miRNAs in parasitic nematodes

The selective enrichment in nematode miRNAs associated with the host AGO1, together with the recurrent deployment of highly conserved miRNA families by phylogenetically distant helminth parasites, supports the view that cross-organism small-RNA communication has evolved as a stereotypical virulence strategy. In addition to identifying the nematode miRNAs loaded into plant AGO1, one of the major implications of this work is the apparently convergent use of ancient miRNA families as secreted signals by different parasitic helminths. For instance, the miR-2, miR-71, and miR-100 families are consistently identified in the EVs of diverse parasitic nematodes or in the serum of their host, highlighting their roles as universal effectors of parasitism (Sotillo et al., 2020; Chowdhury et al., 2024). At family level, miRNA repertoires are broadly conserved, but repeated observations of the preferential release and detection of a subset of highly conserved miRNAs, such as members of the miR-2 family, in host-associated compartments provides strong evidence against passive leakage, instead suggesting that there is positive selection for cross-species functionality. The situation is different for the highly expressed secreted miR-100 and miR-71. Importantly, convergence does not necessarily imply a conservation of precisely the same target genes. Instead, parasites may repeatedly exploit a limited number of host regulatory hubs, such as that for immunity, through the use of conserved target sites or different species-specific sites with similar physiological outputs. Our data therefore support a model in which cross-kingdom miRNA delivery constitutes a recurrent, evolutionarily reinforced virulence strategy. One limitation of our AGO1-RIP dataset concerns let-7. We excluded AGO1-associated min-let-7 as likely contamination because it was also detected in the negative control (non-infected roots incubated with free-living J2s). However, let-7 has been repeatedly reported in extracellular vesicles/secretions and host-associated fluids in multiple parasitic systems, including plant-parasitic nematodes (Sotillo et al., 2020; Ste-Croix et al., 2023), and it co-localizes with the secreted miR-100 in a conserved genomic cluster in M. incognita. Thus, although we opted for a conservative interpretation here, we cannot rule out that let-7 is genuinely secreted and loads into host AGO1 during infection; resolving this will require further analysis.

Together, our data support a working model in which M. incognita implements a highly selective, miRNA-driven cross-kingdom RNAi program during infection. A restricted subset of nematode miRNAs is preferentially sorted for secretion independently of their abundance in the parasite, delivered into root giant feeding cells, and selectively loaded into host AGO1, positioning them within the core plant silencing machinery. A fraction of these AGO1-bound miRNAs then converges on key host regulatory hubs—hormone-mediated immunity, metabolic reprogramming, and cellular organization/cytoskeletal dynamics—thereby coordinating broad transcriptional and physiological outputs that attenuate defense while promoting giant-cell development and sustained feeding-site function across hosts. Consistent with this model, miR-2b emerges as an effector that enhances giant feeding cell formation/maintenance in infected roots. Future work should define the *in vivo* contribution of individual secreted miRNAs, elucidate the mechanisms underlying selective sorting and delivery, and extend analyses to other transferred nucleic acids to capture the full scope of cross-organism molecular exchange during plant–nematode interactions.

## METHODS

### Plant Growth Conditions and Inoculation

*Solanum lycopersicum* (cultivar M82) seeds were sown on 0.8% agar plates, and incubated at 22°C in darkness for four days. Seedlings were transplanted into a 1:1 mixture of soil and sand (B5) and grown in a controlled-climate chamber under a 16-hour light/8-hour dark photoperiod, at 22°C and 60% relative humidity.

*Arabidopsis thaliana* (ecotype Col-0) seeds were stratified for 48 hours at 4°C, sown in potting soil and placed in a germination chamber for one week (8-hour light/16-hour dark photoperiod, 22°C, 80% relative humidity). Individual seedlings were transplanted into single soil pots and placed in a growth chamber under an 8-hour light/16-hour dark cycle at 22°C and 60% relative humidity.

One month after germination, each plant was inoculated with second-stage juveniles (J2) of *Meloidogyne incognita* (strain Morelos), 500 such larvae for *Arabidopsis* and 1500 for *S. lycopersicum*. Fourteen days post-infection, galls harvested from infected plants or roots from uninfected plants (as a control), were flash-frozen in liquid nitrogen, and stored at -80°C for further analysis.

### RNA extraction and sequencing

Preparasitic J2 were flash frozen in liquid nitrogen and ground to a powder in a tissue lyser (Retsch; MM301) with 4 mm tungsten balls at a frequency of 30 Hertz for 30 seconds. Total RNA was extracted from these samples with the miRNeasy Mini Kit (Qiagen), according to the manufacturer’s instructions, but with three additional washes in RPE buffer.

The quality of the small RNAs generated by AGO1-RIP and of the total J2 RNA were checked with a Bioanalyzer High-Sensitivity sRNA kit High-Sensitivity RNA kit respectivelu. Small RNAs and total J2 RNAs were sent to the Beijing Sequencing Institute (Hong Kong, China) for the construction of libraries and sequencing. The full raw sequencing data were submitted to the GEO database (http://www.ncbi.nlm.nih.gov/geo/).

### M. incognita MiRNA de novo prediction

Two complementary approaches were used for small RNA prediction in the nematode, to increase robustness. First, miRNAs were predicted with miRDeep2 (v2.0.0.8), which was run independently on each sequencing library (Friedländer et al., 2012). Only miRNA candidates detected in at least two independent libraries were retained for downstream analyses. In parallel, miRNA prediction was also performed with ShortStack (v3.8.5) (Axtell, 2013). We used a multi-step strategy to decrease the false-positive rate: (i) candidate loci were defined from the initial ShortStack predictions; (ii) loci were compared across libraries with MultiIntersectBed, retaining only those supported by at least three libraries; and (iii) the retained loci were merged to generate a consolidated locus set. ShortStack was then rerun independently on each library, using this consolidated set, to refine miRNA calls. Finally, miRDeep2- and ShortStack-derived predictions were compared and merged with the aid of a custom Perl script, yielding a final high-confidence set of predicted miRNAs.

### Co-immunoprecipitation of AGO1-Associated sRNAs

Frozen galls were harvested and ground into a fine powder in liquid nitrogen: 5 g from galls of *S. lycopersicum* and 1 g from galls of *Arabidopsis*. We mixed 5 g of root tissue from uninfected tomato plants with *M. incognita* J2 larvae (1000 J2 per gram of fresh uninfected root) as a control. AGO1 immunopurification was performed as described by Dunker et al., (2021) with a slight modification. Non-specific co-purification was minimized by incubating the extract with Protein A agarose beads (Roche) for 30 min before washing. The beads were removed by centrifugation at 300 x *g* for 1 min at 4°C. AGO1 immunoprecipitation was performed by incubating the cleared extract with 1 µg per gram of tissue on a rotating wheel for 4 hours at 4°C. We added Protein A agarose beads at a concentration of 100 µL per 5 mL IP buffer, and the mixture was incubated overnight at 4°C with continuous rotation.

### AGO1-associated small-RNA sequencing analysis

AGO1-immunoprecipitated (AGO1-IP) sRNA libraries were processed with a reproducible bioinformatics pipeline, starting from raw FASTQ files, as follows

i. Read preprocessing and rRNA depletion: rRNA contaminants were removed by filtering raw reads with SortMeRNA v4.3.6 against the SILVA rRNA database (v138.1) with default parameters (Martinez et al., 2008). The remaining reads were size-selected to retain only sequences of between 18 and 25 nt. We ensured that rRNA depletion was exhaustive by performing a second specific filtering step with Bowtie v1.3.1 (Langmead et al., 2009). Reads were aligned against the SILVA 138.1 database with stringent parameters adapted for short sequences (-l 10 -e 70 -v 0, allowing only perfect matches) (Quast et al., 2013). The non-aligning reads (rRNA-free) were extracted from SAM files with Samtools v1.14 (bam2fq) for downstream analysis (Danecek et al., 2021).
ii. Genome mapping and quantification: The rRNA-free sRNA reads were mapped onto the reference genomes of *Solanum lycopersicum* (Heinz1706 -v3.00), *Meloidogyne incognita* (v3), and a combined tomato-nematode reference genome with Bowtie v1.3.1. We ensured computational reproducibility by performing all alignments in a Singularity container environment. The following alignment parameters were applied: perfect matches only (-v 0), reporting all valid alignments (-a), and retaining only the best alignments (--best --strata). Aligned sequences were extracted and collapsed into unique tags. The abundance of each unique sequence was quantified across conditions and reference genomes with custom Perl scripts.
iii. miRNA homology search and annotation: Homology-based annotation was performed on unique sequences with HMMER v3.4 (nhmmer) in DNA mode (Wheeler and Eddy, 2013). Search targets included miRBase, and sets of predicted miRNAs for both *S. lycopersicum* and *M. incognita* (Kozomara et al., 2019). Alignments were optimized for short reads with a score threshold of -T 5 and the F3 filter enabled. Significant hits were retained to generate the final annotation of AGO1-associated sRNAs.
iv. Thresholds of 50 reads per million present in two replicate galls for each sequence were applied and the final list of sRNAs was manually curated with the retention of perfect alignments against the Rfam and Ensembl databases to eliminate all sequences corresponding to tRNA, snoRNA, snRNA, long non coding RNAs (lncRNAs) and all sense mRNAs.

### Degradome analysis

The degradome sequencing of tomato galls (Noureddine et al., 2023) was analysed with CleaveLand 4.0 (Addo-Quaye et al., 2009), using the default parameters and the secreted nematode miRNAs. *In sillico* prediction was performed with psRNATarget (default Schema V2) (Dai et al., 2018), using the list of secreted miRNA and the *Solanum lycopersicum* cDNA library ITAG3.0 as input.

### Quantification of miRNA Cleavage Activity With the Dual-Luciferase Reporter

DLR assays were performed with the DLR Assay System (Promega). For each plant target/nematode amiRNA pair, two leaves from four-week-old *Nicotiana benthamiana* plants were infiltrated with an *Agrobacterium* mixture containing pK2GW7-amiRNA and pGrDL-TS constructs at a 1:5 ratio in infiltration buffer (2 mM MgCl, 2 mM MES, 100 µg/mL acetosyringone). The mixture was incubated for 4 hours before infiltration. The leaves were harvested 72 hours after infiltration, ground into a fine powder in liquid nitrogen with 4 mm tungsten beads in n MM400 vibratory crusher (Retsch; MM30) and stored at -80°C. Five mg of ground leaf tissue was resuspended in 250 µL 1X Passive Lysis Buffer (PLB; Promega). A 15 µL aliquot of the lysate was diluted in 360 µL buffer for analysis. *Renilla* and firefly luciferase activities were measured with a SAFAS luminometer. Measurements were performed on 96-well plates with 15 µL of leaf extract per well. Firefly activity was measured 10 seconds after the injection of 75 µL Luciferase Assay Buffer II (Promega), and then 10 seconds after the addition of 75 µL Stop & Glow reagent for *Renilla* luciferase activity (Promega). The mean luminescence ratio over 10 seconds was calculated for each luciferase in the eight biological replicates of each nematode miRNA/tomato target-site pair, with at least two independent experimental replicates. Ratios were normalized against those obtained with the pGrDL plasmid lacking a target site and are presented as log values. Statistical comparisons of F-Luc/R-Luc ratio normalized against pGrDL_SPb and converted into log_2_ ratios between each pGrDL_Target-Site and the pGrDL_SPb control co-infiltrated with the same amiRNA were performed with unpaired Student’s *t*-tests in GraphPad Prism v8.

### Microsynteny analysis

We used MCScanX (Wang et al., 2012) with default parameters to determine whether the *MIR* regions of genus *Meloidogyne* genus displayed conserved synteny. BLASTp files and GFF3 annotations were provided for all the available *Meloidogyne* species (*M. graminicola*, *M. chitwoodi*, *M. hapla*, *M. enterolobii*, *M. arenaria*, *M. javanica*, *M. incognita* and *M. luci*) and the closest outgroup species, *P. penetrans*. MCScanX was ran with the default parameters. However, the -b interspecies parameter was used to identify collinear blocks between different species, with an e-value of 0.001, and the -s 4 parameter was used to impose a requirement of at least four genes per collinear block. We obtained a set of syntenic blocks alongside a detailed list of syntenic gene pairs, which we used to identify *MIR* regions in the collinearity file and to determine in which *Meloidogyne* species these regions were conserved.

### *S. lycopersicum* hairy-root production and infection assay

Artificial microRNAs (amiRNAs) were expressed transiently in *S. lycopersicum* by *Agrobacterium rhizogenes*-mediated hairy-root transformation. The binary vector pK2GW7-amiRNA was introduced into *Agrobacterium tumefaciens* strain C58C1 harboring the *A. rhizogenes* pRiA4 plasmid and maintained under chloramphenicol, streptomycin and kanamycin selection pressure. Tomato (cv. M82) seeds were surface sterilized by HCl treatment and germinated on Murashige and Skoog (MS) medium for 10 days. Seedlings were then subjected to stem agroinfiltration by injection of the *Agrobacterium* suspension (OD=1) with a syringe. Ten days post-inoculation, stem segments bearing emerging hairy roots were excised and transferred to MS medium supplemented with 3% sucrose, 0.3% phytagel, 0.05% MES, 500 µg/mL ticarcillin, 200 µg/mL cefotaxime, and 30 µg/mL kanamycin for the selection of transformed roots. Hairy roots carrying the pK2GW7-amiRNA construct were verified by PCR and transferred to MS medium without antibiotics. They were allowed to regenerate for one week and individual hairy roots were then inoculated with 250 freshly hatched preparasitic second-stage juveniles (J2s). The J2s were surface-sterilized with HgCl and treated with streptomycin before inoculation. At 14 days post-inoculation, galls were collected and fixed in 30% formaldehyde in 1× PBS under a vacuum. The tissues were cleared with benzyl alcohol/benzyl benzoate (BABB) and the feeding sites were then measured.

### Benzyl alcohol/benzyl benzoate clearing and feeding site measurement

After one week of regeneration, individual hairy roots were inoculated with 250 freshly hatched sterilized preparasitic J2s. Measurements of giant cell area within 14 dpi galls were performed as previously described (Cabrera *et al*., 2018; Noureddine *et al*. 2023).

## AUTHOR CONTRIBUTIONS

A.D., S.J., and B.F. conceived the study and designed experiments. Unless otherwise specified, A.D. performed all of the experiments and conducted data analysis. MdR performed the bioinformatic analysis. E.S. performed microsynteny analysis. A.W., Y.N. and A.P.C initiate the AGO1-RIP. K.M. genotyped the hairy roots. P.F. performed the BABB experiments. O.Y. and A.T.M. performed some DLR analyses. J.Z. and L. N. supervised some sRNAs analysis. M.Q. take part to the writing. A.D., S.J. and B.F. wrote the article.

## ACKNOWLEDGMENTS

We would like to thank Etienne Danchin and Cei Abreu for their advices on the RKN genome and microRNA analysis, Sophie Mantelin for her advice on cloning, Julien Pirello and Lydie Tessaroto for their help in setting up the tomato root transformation.

## SUPPLEMENTAL INFORMATIONS

- Supplemental Figure 1: Characteristics of *M. incognita* mature and precursor miRNA sequences.
- Supplemental Figure 2: Experimental test of tomato genes identified as predicted targets of nematode miRNAs.
- Supplemental Figure 3: Conservation of the tomato miR-2b target site of SlMT2 in *Solanum lycopersicum* and *Arabidopsis thaliana* methyltransferase-encoding genes.
- Supplemental Figure 4: Genome organization and sequence conservation of the miR-2 family within the phylum Nematoda
- Supplemental Figure 5: MicroRNAs secreted by nematodes, trematodes and cestodes.
- Supplemental Figure 6: Western-blot analysis of SlAGO1-IP
- Supplemental Table 1: *M. incognita MIR* gene prediction
- Supplemental Table 2: Genomic clusters of MIR genes
- Supplemental Table 3: miRNA expression in J2 G7 G14
- Supplemental Table 4: SlAGO1-RIP sequencing
- Supplemental Table 5: Analysis of the differential expression of miRNA in SlAGO1-RIP
- Supplemental Table 6: Tomato target prediction based on degradome analysis and CleaveLand V4 cat ≤2
- Supplemental Table 7: Tomato target prediction with psRNATarget Exp. Score ≤3.5
- Supplemental Table 8: Selected *S. lycopersicum* miRNA targets
- Supplemental Table 9: AtAGO1-RIP sequencing
- Supplemental Table 10: Microsynteny of secreted miRNA in *Meloidogyne* spp.
- Supplemental Table 11: List of primers and oligomers used in this study
- Supplemental Figure 8: Anti-AGO1 western blot on SlAGO1-IP

