## Supplemental Figure 1 for "Plant-parasitic nematode microRNAs hijack plant AGO1 to induce host-cell reprogramming"

### SUPPLEMENTAL FIGURES

- Supplemental Figure 1: Characteristics of *M. incognita* mature and precursor miRNA sequences.
- Supplemental Figure 2: Experimental test of tomato genes identified as predicted targets of nematode miRNAs.
- Supplemental Figure 3: Conservation of the tomato miR-2b target site of SlMT2 in *Solanum lycopersicum* and *Arabidopsis thaliana* methyltransferase-encoding genes.
- Supplemental Figure 4: Genome organization and sequence conservation of the miR-2 family within the phylum Nematoda
- Supplemental Figure 5: MicroRNAs secreted by nematodes, trematodes and cestodes.
- Supplemental Figure 6: Western-blot analysis of SlAGO1-IP

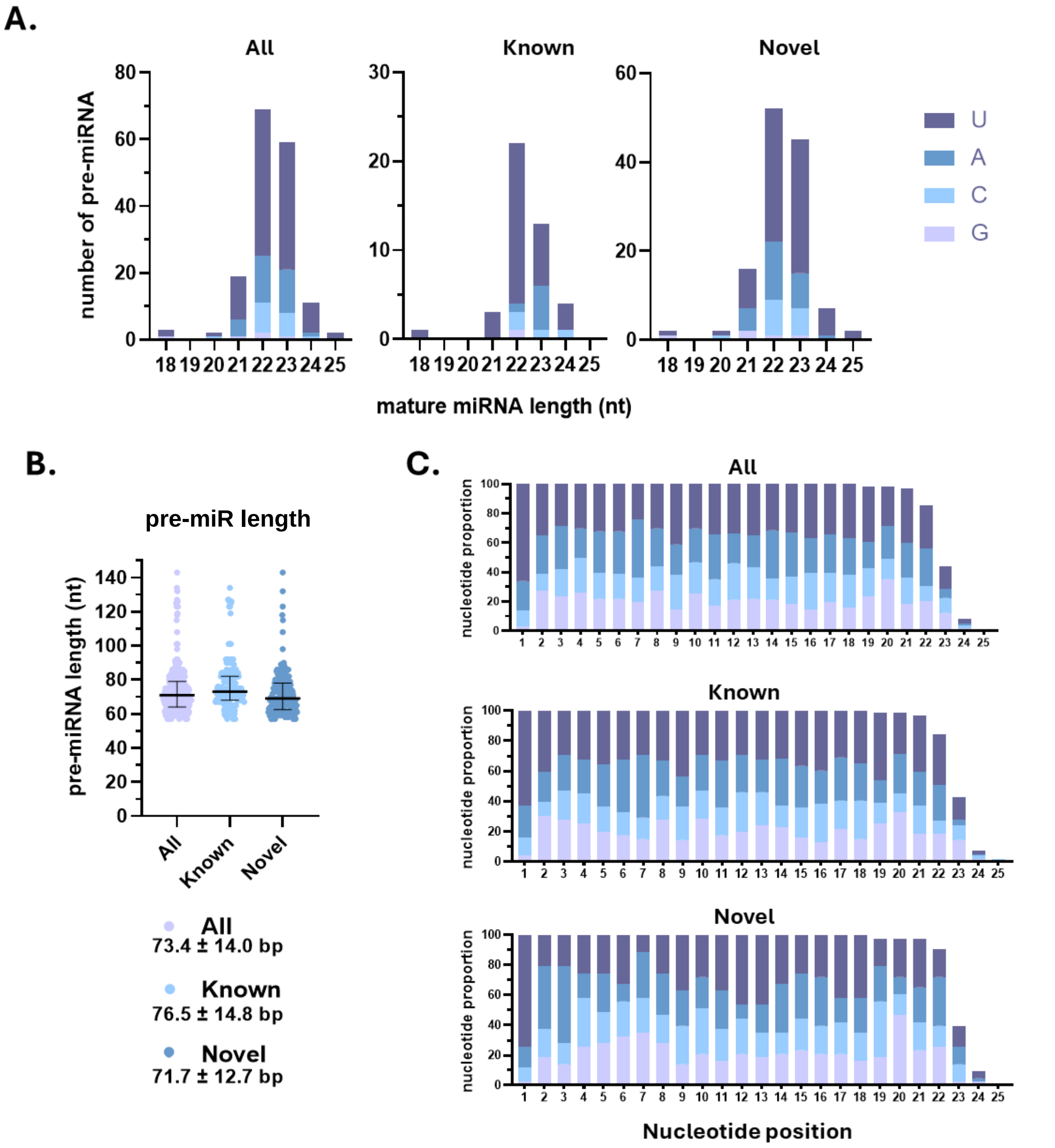


**Supplementary Figure 1. Characteristics of *M. incognita* miRNA mature and precursor sequences. (A)** Profiles of the 5′ nucleotide and length of mature miRNAs, **(B)** length distribution of miRNA precursors and **(C)** Nucleotide composition at each position of mature miRNAs for all identified precursors, including known and novel miRNAs.

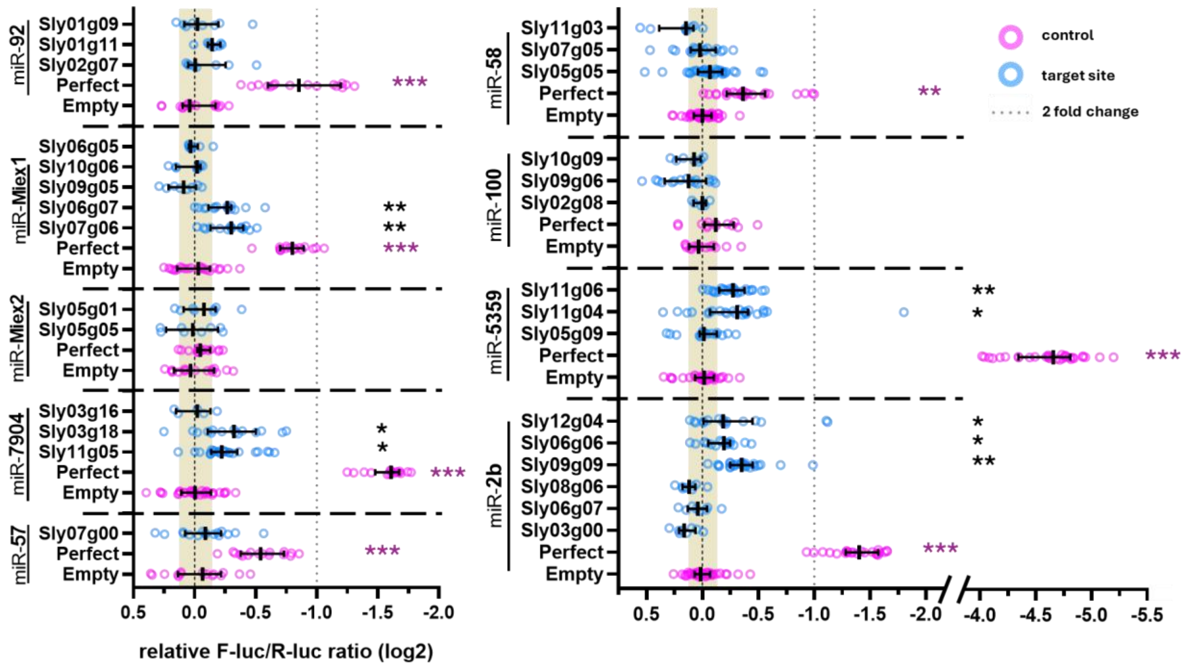


**Supplemental Figure 2 : Experimental test of tomato genes predicted to be targeted by nematode min-miRNAs**. Min-miRNAs and their predicted tomato targets were co-expressed in *N. bethanmiana* leaves to test the silencing efficiency. The predicted target sequence was expressed fused to the firefly lmuciferase coding sequence in a DLR system that enables the constitutive expression of renilla luiferase as an internal control. Relative ratio of Firefly luciferase (F-luc) and Renilla luciferase (R-luc) in log2 measured on *N.benthamiana* leaves co-infiltrated with each couple miRNA/target-site and normalized on negative control. Sequence perfectly complementary to the tested min-miRNAs (perfect) was used as a postitive silencing control. “Spacer” corresponding to 5’UTR natural sequence of Firefly luciferase negative control assays use sequences with little/no complementarity to the tested miRNA. Asterisk represents statistically significant differences using unpaired student t-test : $^{*}P < 0.05$, $^{**}P < 0.001$, and $^{***}P < 0.0001$.

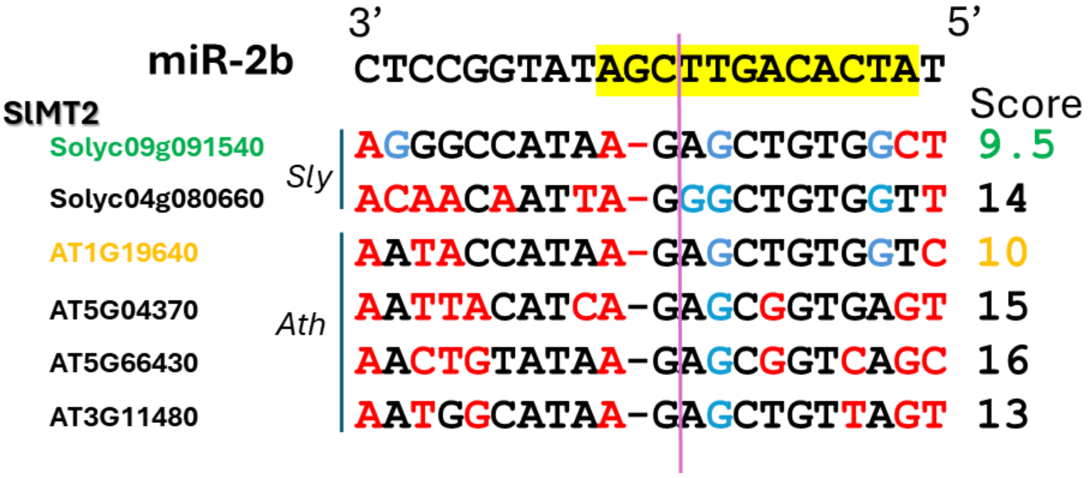


**Supplemental Figure** 3 : **Conservation of the tomato miR-2b target site of SlMT2 in *Solanum lycopersicum* and *Arabidopsis thaliana* Methyl transferases encoding genes**. Paralogs of *S. lycopersicum* (*Sly*) target genes and homologs from *A. thaliana* (*Ath*) were retrieved from Ensembl and aligned to identify potential conserved cleavage sites. Min-miRNA nucleotides highlighted in yellow correspond to positions 2-13, which are considered a key pairing region. Nucleotides shown in red indicate mismatches, while those in blue represent G:U pairings. Alignment scores were calculated using the following pairing parameters: mismatch = 1; G:U pairing = 0.5; gap = 2; scores were doubled when occurring within the yellow-highlighted region. Gene IDs shown in green correspond to targets for which cleavage activity by min-miRNAs was confirmed by dual-luciferase (DLR) assays; those shown in blue correspond to targets predicted by psRNATarget (expectation score < 5); and those shown in yellow correspond to transcripts displaying sufficient complementarity to be considered potential targets, based on a maximum score threshold of 10. The tomato *SlMT2* (*Solyc09g091550*) and the *Arabidopsis AtJMT* (*AT1G19640*) display a conserved target site.

**A** **B**

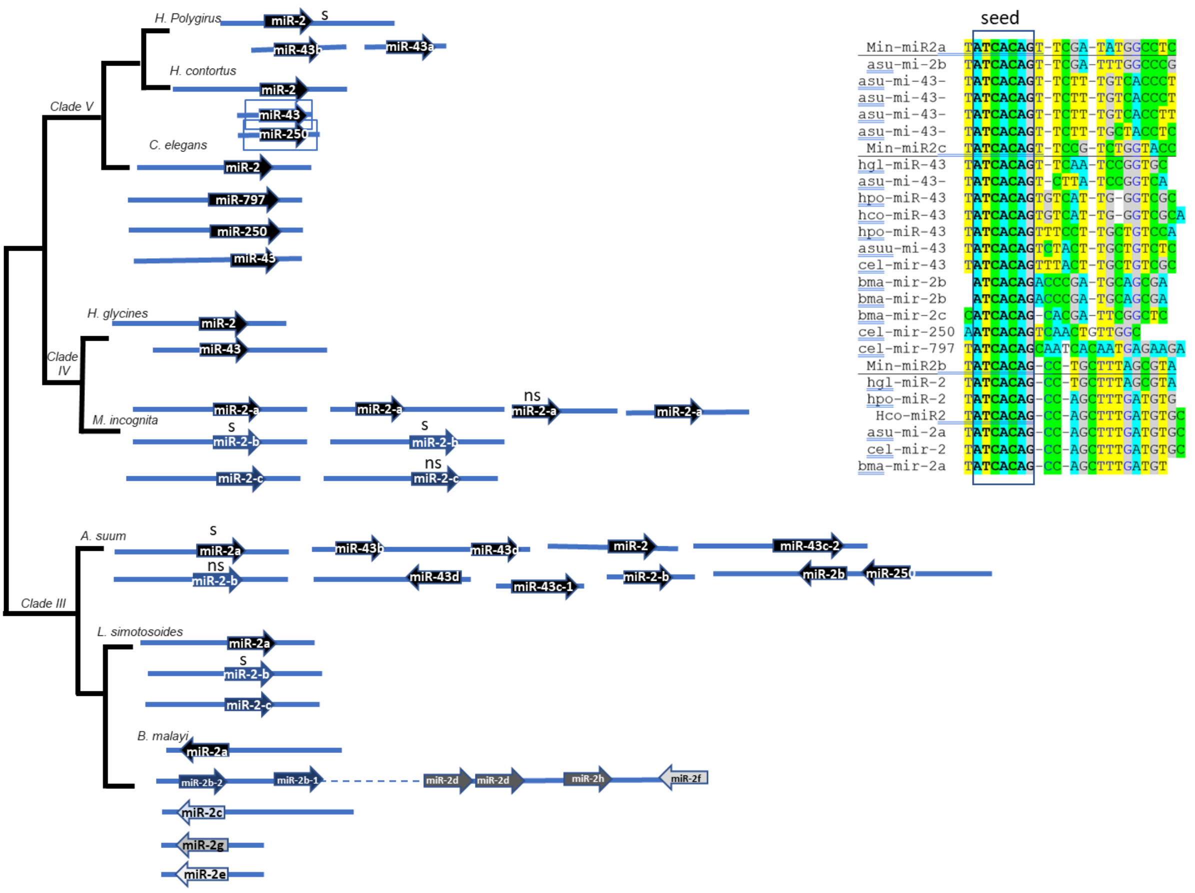


**Supplemental Figure 4: Genome organisation and sequence conservation of the miR-2 family within the phylum Nematoda. (A)** Phylogenetic tree of miR-2 family genes in animal-parasitic nematodes *Ascaris suum*, *Litomosoides sigmodontis*, *Brugia malayi*, *Haemochus contortus* and *Heligmoides polygirus*, in the plant-parasitic nematodes *Meloidogyne incognita* and *Heterodera glycines* and in the free living nematode *Caenorhabditis elegans*. **(B)** Alignment of miR-2mature sequences from nematodes.

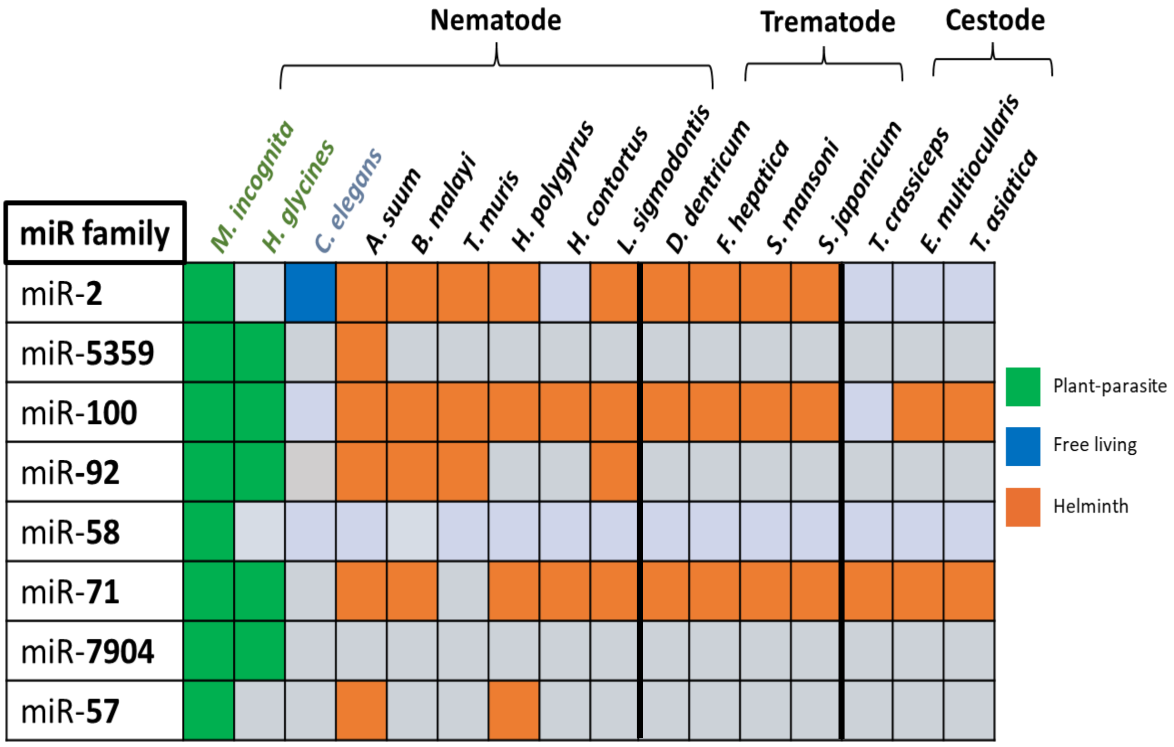


**Supplemental Figure 5: MicroRNAs secreted by nematodes, trematodes and cestodes.** SlAGO1 associated min-miRNAs are found in EVs from several parasitic nematodes and helminth. MicroRNAs were identified in the secretion, in exosomes or in the host tissue or serum of the plant parasitic nematodes *M. incgnita* and *Heterodera glycines*, of the free living nematode *Caenorhabditis elegans*, of the animal parasitic nematodes *Ascaris suum*, *Brugia malayi*, *Trichuris muris*, *Heligmoiides polygirus*, *Haemonchus contortus*, *Litomosoides sigmodontis*, of the animal parasitic trematodes Dicrocoelium dendriticum, Fasciola hepatica, *Schistosoma mansoni and japonicum* and of the animal parasitic cestodes *Taenia crassiceps*, *Echinochoccus multilocularis* and *T. asiatica* (Sotillo *et al*. 2020; Ste croix *et al*. 2023)

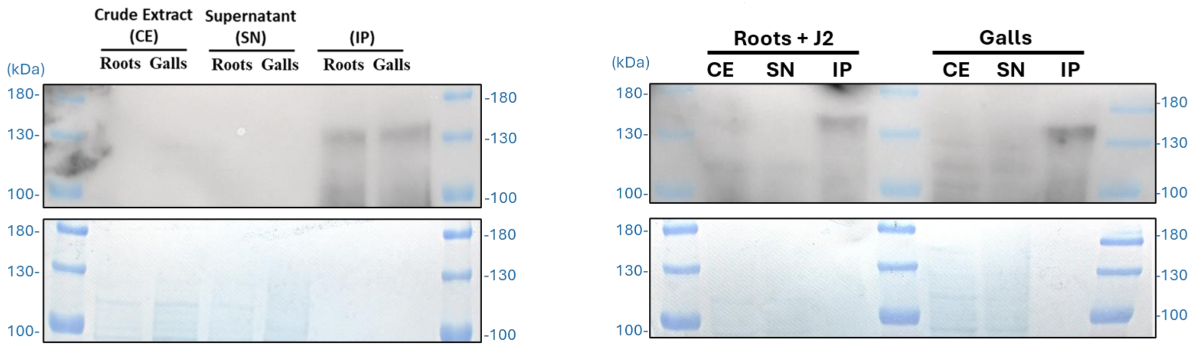


**Supplemental Figure 6: Western blot analysis of SlAGO1-IP.** SDS–PAGE membranes from SlAGO1-RIP samples prior to RNA extraction, visualized using a Vilber imaging system based on the anti-AGO1–HRP signal (top) and total proteins stained with Coomassie Blue (bottom). CE = crude extract; SN = supernatant; IP = immunoprecipitation.
